# Composition of minor ampullate silk makes its properties different from those of major ampullate silk

**DOI:** 10.1101/2022.12.12.520175

**Authors:** Hiroyuki Nakamura, Nobuaki Kono, Masaru Mori, Hiroyasu Masunaga, Keiji Numata, Kazuharu Arakawa

## Abstract

Spider’s minor ampullate silk, or MI-silk, exhibits distinct mechanical properties and water resistance compared to its major ampullate counterpart (MA-silk). The principal protein constituent of MI-silk is known as minor ampullate spidroin, or MiSp, and while its sequence has been deciphered and is thought to underlie the differences in properties with MA-silk, the composition of MI-silk and the relationship between its composition and properties remain elusive. In this study, we set out to investigate the mechanical properties, water resistance, and proteome of MA-silk and MI-silk from *Araneus ventricosus* and *Trichonephila clavata*. We also synthesized artificial fibers from major ampullate spidroin, MaSp1 and 2, and MiSp to compare their properties. Our proteomic analysis reveals that the MI-silk of both araneids is composed of MiSp, MaSp1, and spidroin constituting elements (SpiCEs). The absence of MaSp2 in the MI-silk proteome and the comparison of the water resistance of artificial fibers suggest that the presence of MaSp2 is the reason for the disparity in water resistance between MI-silk and MA-silk.

## INTRODUCTION

Spiders use up to seven different silks for various purposes throughout their life stages, including use as a dragline, foraging, nesting, communication, and to protect the eggs.^1^ These silks have different morphologies and mechanical properties tailored for their ecological use.^2^ Each silk type has a corresponding silk gland and associated spigots. These silk glands are also morphologically different from each other, as their secretion comprises different amino acid compositions, suggesting differing constituents.^3,4^

Major ampullate silk (MA-silk) and minor ampullate silk (MI-silk) are two types of silks produced by major and minor ampullate glands. MA-silk is used as the dragline or in the construction of the radii, frame and anchor lines of orb webs.^1,5^ MA-silk is known for its substantial toughness, combining high tensile strength and high elasticity.^5,6^ The molecular structure of MA-silk is modeled as a semicrystalline material composed of recurrent beta-pleated sheet crystallites and amorphous domains.^7^ Nuclear magnetic resonance (NMR) and wide angle X-ray scattering (WAXS) confirmed the model and assigned the beta-pleated sheet crystallite for poly alanine.^8–10^ Components of MA-silk are thought to be major ampullate spidroin 1 (MaSp1) ^11^ and major ampullate spidroin 2 (MaSp2) ^12^, which have Gly-Pro and Glu-Glu motifs in their amino acid sequences.^13^ Recently, another type of spidroin named MaSp3 was identified in several species using target capture sequencing.^14,15^ Studies utilizing a multiomics approach revealed that MaSp3 is one of the main components of MA-silk.^16,17^

MI-silk, a comparably thinner fiber than MA-silk, is often used to construct the temporally (auxiliary) spiral, which is used during orb web building as a walking scaffold^18^ and as a bridging line when reaching distant places.^19^ Mattina et al. found using proteomic analysis that *Latrodectus hesperus* uses MI-silk as a constituent of prey-wrapping silk.^20^ Several lineage-specific uses are also observed, such as *Araneus diadematus* using a MA-silk and MI-silk pair to construct most radii in their orb web.^21^ In many species of Araneoidea, temporary spirals are reported to be cut out and removed during the construction of capture spirals;^22^ however, Nephilinae spiders do not remove this temporary spiral and retain it in their finished orb webs. Hesselberg and Vollrath reported that the mechanical properties of temporary spirals in Nephilinae spider’s orb webs are identical to those of MA-silk, suggesting that the temporary spiral of Nephilinae spider’s orb webs might not consist of MI-silk^23^. The ecological implications of the use of MI-silk therefore remain largely elusive, but an economical preference in applications not requiring high tensile strength is suggested due to the reduced amount of protein by smaller diameter of the MI-silk.^1^

The mechanical property of MI-silk is clearly different from that of MA-silk. It has higher elasticity and lower Young’s modulus and tensile strength than MA-silk,^2^ while the molecular structure of MI-silk is also a semicrystalline structure, and the crystallite is a β sheet of poly alanine.^24^ A known component of MI-silk is minor ampullate spidroin (MiSp), for which Colgin and Lewis reported two paralogs, namely, MiSp1 and MiSp2 from the *Trichonephila clavipes* minor ampullate gland.^25^ Both MiSp1 and MiSp2 have GGX motifs as well as GA repeats in the amorphous domains and poly alanine in crystalline regions. MiSps often also contain nonrepetitive Ser-rich spacer regions of approximately 140 residues in length, which was suggested in a study utilizing partial recombinant protein to be involved in fiber assembly under the existence of shear force.^26^

The most significant difference between MA-silk and MI-silk is their reaction to water molecules. When MA-silk is exposed to water, it undergoes supercontraction up to approximately 50% of its original length. In 1977, work reported that MA-silk undergoes supercontraction when wetted, but MI-silk shows smaller contraction than MA-silk.^27^ In their comprehensive study of MA-silk and MI-silk, Guinea et al. confirmed that MI-silk does not have the ground state of MA-silk, and permanent deformation occurs after contraction.^28^ The mechanical properties of MA-silk are significantly modulated after supercontraction, where the strain at break is increased and the tensile strength is decreased. The change is also known to be correlated with its capacity to shrink;^29^ accordingly, structural analysis with WAXS and Raman spectroscopy suggest that oriented β-sheet nanocrystallites and oriented amorphous components in MA-silk lose their orientation upon supercontraction.^30–32^ The loss of orientation is larger for MA-silk than for MI-silk,^31^ mirroring the propensity for supercontraction. Several amino acid sequence motifs are suggested to be responsible for supercontraction; namely, YGGLGS(N)QGAGR of MaSp1,^33^ GPGXX of MaSp2^34^ and GXG in MaSp1^35^.

Quantitative and comprehensive compositional analysis of protein constituents of the spider silks is beginning to be uncovered for MA-silk, with high-quality genome data coupled with high-sensitivity mass spectrometry. Kono et al. reported that MA-silk contains approximately 1∼5% spider-silk constituting element (SpiCE), along with various spidroin paralogs other than MaSp1 or MaSp2.^16,17^ Experiments *in vitro* with recombinant SpiCE proteins suggested that the addition of a minute (1%) amount of SpiCE to artificial silk significantly boosts its mechanical properties^16^. Early works on the amino acid composition of minor ampullate gland secretions were first reported by Andersen, who found that it had nearly the same composition as fibroin from the silk of *Bombyx mori*^3^. Work and Young performed forcible silking to separate MA-silk and MI-silk using a microscope and successfully analyzed the amino acid composition of both silks. Their observation showed that the content of Proline was significantly lower in MI-silk, and the variability of amino acid composition between species was lower in MI-silk than MA-silk^36^. Chaw et al. reported protemic analysis of MA and MI glands in *Latrodectus hesperus*, where they successfully identified multiple silk-gland specific transcripts, but the contribution of these components to the mechanical properties of the silks wass yet to be explored^14^.

To this end, we established separate silking of MA-silk and MI-silk and used this method to observe differences in the mechanical properties and proteomic compositions of MA-silk and MI-silk of *Araneus ventricosus*. Dragline silk often contains both MA-silk and MI-silk; therefore, we retrieved these silks separately under a microscope. We have especially focused on supercontraction, observing the change in mechanical properties in a time series, where we performed tensile testing 7.5 h, 24 h and 48 h after immersion until silks completely dried. Moreover, we analyzed the protein composition of MA-silk and MI-silk of *A. ventricosus* and *Trichonephila clavata* and analyzed the contribution of MiSp protein using recombinant silk to examine component-function relationships.

## Experimental Section

### Separate forcible silking

Mature *Trichonephila clavata* females were collected in Nagoya city, Aichi, Japan in October 2019 and Tsuruoka city, Yamagata, Japan in November 2021. Mature *Araneus ventricosus* females originally captured in Tsuruoka city, Yamagata, Japan in September 2018 were bred and reared in the laboratory. Spiders were fed a cricket twice a week. Collection of MA-silk and MI-silk samples was conducted with the forcible silking method^36–38^. Spiders were gently immobilized using two pieces of sponge and locked with rubber bands and then placed under a stereomicroscope. After immobilization, silks from the targeted spinnerets were obtained by tweezers and attached to the end of the bobbin. In this step, to ensure the reeling of MA-silk and MI-silk separately, we observed spinnerets with a stereomicroscope to identify silks while reeling. To prevent contamination, we fixed either of the silks with pieces of masking tape and forcibly silked only one type at a time. For example, when we silked MA-silk from one spinneret, we fixed a pair of MI-silks, and vice versa. Silking was constantly monitored under the microscope, confirming the glandular origin of silks (Figure 1A). Dedicated reeling devices were used for silking with a constant reeling speed (1.28 m/min). Forcible silking was continued for 30 to 60 min. After forcible silking, the bobbin-reeled silk was put in a plastic bag and stored in a dry box (relative humidity was kept lower than 30%) at room temperature.

**Figure 1.**
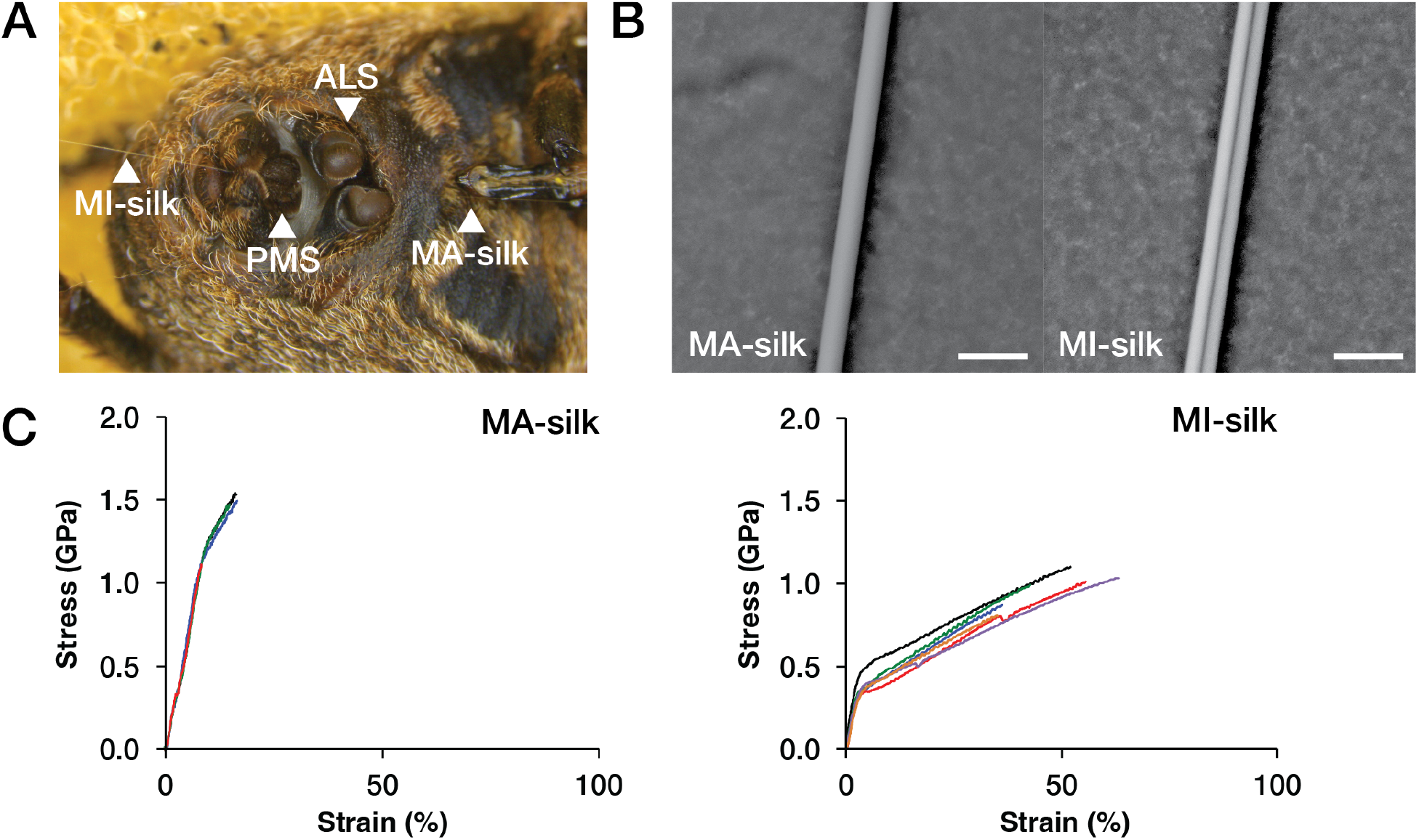
Forcible silking of *A. ventricosus* MA-silk and MI-silk, morphology, and stress‒strain curve of silks

### Maximum supercontraction

The supercontraction of MA-silk and MI-silk was evaluated according to a previous method.^28,37^ Test pieces were prepared by cutting fragments of 5 to 10 cm (L_0_) from MA-silk and MI-silks reeled from *A. ventricosus*. Small pieces of masking tape were affixed on either end of the test pieces. The fibers were immersed in RO (reverse osmosis) water for 1 min to allow contraction. The length of fibers was measured (L_1_) immediately after removal of water, and the percentage contraction was calculated by immersion in water as (L_0_ – L_1_)/L_0_ * 100 and then allowed to air dry for 24 hours in an unrestrained state. The final length of the fiber (L_f_) was measured, and the percentage total contraction after drying was calculated as (L_0_ – L_f_)/L_0_ * 100. Five replicates were performed for each sample. Throughout the experiment, the room temperature was 25°C and relative humidity was around 20%.

### Observation of drying effect of MA-silk and MI-silk during water immersion and subsequent drying

To observe the transition of mechanical properties of contracted MA-silk and MI-silk fibers during drying after water immersion, we measured the mechanical properties of drying fibers intermittently. First, reeled MA-silk and MI-silk fibers were immersed in water. After immersion, the fibers were dried at 20 °C and 20% relative humidity in an unrestrained state. After 2.5 hours, 7 hours and 48 hours of immersion, the mechanical properties and morphology of the fibers were analyzed with tensile tests and scanning electron microscopy (SEM). For each time point, we used at least 3 test pieces.

### Tensile testing

The tensile properties of the fibers were measured using an EZ-S universal tester (Shimadzu, Kyoto, Japan) with a 1 N load cell in an environmental chamber kept at the same relative humidity as other experimental conditions (approximately 20%). The initial length of the single silk fiber was set to 10 mm. The extension speed was applied at 10 mm/min, and the force during testing was measured with a 1 N load cell. Tensile strength, Young’s modulus, strain at break, and toughness were obtained from the resultant stress‒strain curves. For the calculation of tensile strength, the cross-section areas of the fiber samples were calculated based on the diameters determined by SEM observations. Analysis of force‒displacement data and calculation of mechanical properties were conducted as previously described ^38^. Significant digits of mechanical properties were determined in accordance with a previous report ^37^.

### SEM observation and cross-sectional area calculation

The surface morphology and cross-section of the dragline silk fibers were assessed using SEM (Phenom ProX, Thermo Fisher Scientific, Phenom-World BV, Eindhoven, The Netherlands) at V = 15 kV. Samples were mounted on an aluminum stub with conductive tape. The cross-sectional area was calculated assuming a circular cross section. The fiber diameters for calculating the cross-sectional area were measured on each micrograph using ImageJ (NIH, Bethesda, MD, USA).

### Birefringence measurement

The retardation provided by the silk fiber was measured with a birefringence measurement system WPA-100 (Photonic Lattice Inc. Miyagi, Japan) and was analyzed with WPA-VIEW (version 1.05) software, in accordance with a previous method. The birefringence of the dragline silk fiber was calculated from the retardation and silk fiber diameter, which was determined via SEM.

### Wide angle X-ray scattering measurement

Synchrotron WAXS measurements were conducted at the BL45XU beamline of Spring-8, Harima, Japan, in accordance with a previous report.^39^ The X-ray energy was 12.4 keV at a wavelength of 0.1 nm. The sample-to-detector distance for the WAXS measurements was approximately 257 mm. The exposure time for each diffraction pattern was 10 s. The resultant data were converted into one-dimensional radial integration profiles using Fit2D software.^40^ The resultant data were corrected by subtracting the background scattering.

### Simultaneous wide-angle X-ray scattering-tensile test

A sample stretching apparatus (Sentech, Osaka, Japan) was placed in the experimental hutch and controlled outside of the hutch at the BL45XU beamline of Spring-8. The dragline silk fibers were attached to the stretching apparatus. The initial length of the fiber bundle between the fixtures was 13 mm. The strain rate applied to the fibers was 3.3 × 10^−1^ s^-1^. The exposure time for each scattering pattern was 0.099 s.

### Expression and purification of recombinant spidroins

A MiSp recombinant protein with optimized sequences based on *Trichonephila clavipes* was used for the artificial silk. Targeting a size of approximately 50 kDa, we reduced the number of repetitive units of *T. clavipes* MiSp1A1 to seven while keeping the N- and C-terminal domains. The mini-spidroin gene is composed of a 6x His tag (MHHHHHH), a linker (SSGSS), the N-terminal domain, repetitive units, and the C-terminal domain. The MiSp recombinant protein was produced and purified as described previously.^16^ MaSp1 and MaSp2 mini-spidroin lyophilized powders were prepared as described previously.^16^

### Production of fibers from recombinant spidroins

Lyophilized recombinant MiSp, MaSp1 and MaSp2 proteins of 15% (w/w) were dissolved in DMSO and 2 M LiCl and stirred for 30 min at 80 °C. The dope was extruded by a N2 pump with a D = 0.1 mm needle at 80 °C and spun directly into the first coagulation bath comprising MeOH. Fibers were coagulated with a second coagulation bath also comprising MeOH and then washed and stretched in a water bath up to 6x.

### Proteome analysis

Sample preparation for proteome analysis of MA-silk and MI-silk was performed as previously described.^41^ Briefly, reeled silk samples were immersed in 6 M guanidine-HCl buffer (pH 8.5) for MA-silk and 9 M LiBr for MI-silk, followed by freezing and freezing in liquid nitrogen. The samples were thawed and subjected to sonication using a Bioruptor II (BM Equipment Co., Ltd.) for protein extraction. After centrifugation at 15,000 x g for 10 min, the protein concentration of the supernatants was quantified using a Pierce BCA Protein Assay Kit (Thermo Scientific). The extracts containing 10‒50 µg of protein were reacted with DTT for 30 min at 37 °C followed by iodoacetamide for 30 min at 37 °C in the dark. After 5-fold dilution with 50 mM ammonium bicarbonate, the proteins in the samples were digested with chymotrypsin for 16 h at 37 °C. The enzymatic digests were acidified by trifluoracetic acid and desalted using C18 StageTips.^42^ Nano liquid chromatography tandem mass spectrometry (nanoLC‒MS/MS) was performed using a nanoElute ultrahigh-performance LC apparatus and a timsTOFPro mass spectrometer (Bruker Daltonics, Bremen, Germany). Each digested sample was injected into a spray needle column (ACQUITY UPLC BEH C18, 1.7 µm, Waters, Milford, MA, 75 µm i.d. × 250 mm) and separated by linear gradient elution with two mobile phases, A (0.1% formic acid in water) and B (0.1% formic acid in acetonitrile), at a flow rate of 280 µL/min. The composition of mobile phase B was increased from 2% to 35% in 100 min, changed from 35% to 80% in 10 min and kept at 80% for 10 min. The separated peptides were ionized at 1,600 V and analyzed by parallel accumulation serial fragmentationscan.^43^ Precursor ions were selected from the 12 most intense ions in a survey scan (precursor ion charge: 0‒5, intensity threshold: 1,250, target intensity: 10,000).

De novo sequencing and database searches were performed with an error tolerance of 20 ppm for precursor ions and 0.05 Da for fragment ions using PEAKS X+ software (version 10.5).^44^ The protein sequences generated from our draft genome dataset^16,17^ (GCA_013235015.1 for *A. ventricosus* and GCA_019973975.1 for *T. clavata*) were used as a reference database. The abundance of proteins identified from each sample was estimated using the intensity-based absolute quantification (iBAQ) method.^45^ The LC‒MS raw data and the associated files were then deposited in the ProteomeXchange Consortium (accession number: PXDXXXXXX) via the jPOST partner repository (accession number: JPSTXXXXXX).

## Results

### Separate forcible silking of MA-silk and MI-silk

We collected MA-silk and MI-silk from *A. ventricosus* with separate forcible silking. The diameters of MA-silk and MI-silk observed with SEM were 3.40 ± 0.17 µm and 2.11 ± 0.10 µm, respectively (Figure 1B). The diameters were consistent with previous reports of forcibly silked MA-silk and MI-silk.^28^ The stress‒strain curves of MA-silk and MI-silk obtained by forcible silking from *A. ventricosus* were tested in air (Figure 1C). The mechanical properties of the stress‒strain curves are summarized in Table 1. MA-silk has shown higher strength and lower extensibility than MI-silk. The Young’s modulus of each silk was almost the same. The trend of mechanical properties of MA-silk and MI-silk is consistent with earlier reports of *Argiope* and *Trichonephila*.^2,28^

**Table 1.** Percentage of contraction of MA-silk and MI-silk.

|  | MA-silk | MI-silk |
| --- | --- | --- |
| % contraction by immersion in water | 51.2 ± 2.6 | 6.9 ± 1.7 |
| % total contraction after drying | 51.6 ± 2.5 | 6.9 ± 1.7 |

### Wide angle X-ray scattering of MA-silk and MI-silk

Comparison of the nanostructure of MA-silk and MI-silk of *A. ventricosus* was conducted with synchrotron wide-angle X-ray scattering (WAXS). By subtracting the MI-silk scattering profile from MA-silk, the difference in crystalline state was characterized (Figure S1). MA-silk’s crystallite size of the *a*-axis and *b*-axis was larger than MI-silk, and the sizes of the *c*-axis were similar for both. In a previous study, *T. clavipes* MI-silk was reported to be more crystalline and to have larger beta-sheet crystallites than MA-silk.^31^ The opposite sizes of MA-silk and MI-silk of *A. ventricosus* with *T. clavipes* might suggest that MI-silks of different species have different structural properties. Then, we observed how the crystal structures of MA-silk and MI-silk deformed while drawing fibers by performing WAXS while drawing (Figure S2). Compared to MA-silk crystallites, MI-silk crystallites were observed to be more prone to lattice deformation. The difference in lattice deformation propensity might be related to the difference in mechanical properties.

### Mechanical properties of MA-silk and MI-silk of *A. ventricosus* before and after immersion in water

We tested the supercontraction of *A. ventricosus* MA-silk and MI-silk to compare the reaction to water of each type of silk. To test supercontraction, we immersed silk samples in water and then measured length immediately. After complete drying (24 hours later), we measured the length of each sample again. The ratio of the length contracted per initial length was defined as the ratio of contraction. Table 1 summarizes the percentage of contraction after immersion in water and subsequent drying. As previous reports described,^27,28^ MA-silk was more contracted than MI-silk. In accordance with previous reports, the total percentages of contraction of MA-silk and MI-silk of *A. ventricosus* were closer to *Araneus diadematus* ^27^ and *Argiope trifasciata* than *Trichonephila inaurata*^28^. *Trichonephila* MI-silk showed higher percentage of contraction than *Araneus* and *Argiope*. Then, we observed deformation of stress-strain curve by water immersion in MA-silk and MI-silk and reversibility thereof. We thus immersed *A. ventricosus* MA-silk and MI-silk into water and measured their mechanical properties after 2.5 hours, 7 hours and 48 hours of immersion with tensile testing. As shown in Figure 2, supercontraction deformed the stress‒strain curve of MA-silk after immersion in water, whereas the stress‒strain curve of MI-silk after immersion in water showed relatively low deformation.

**Figure 2.**
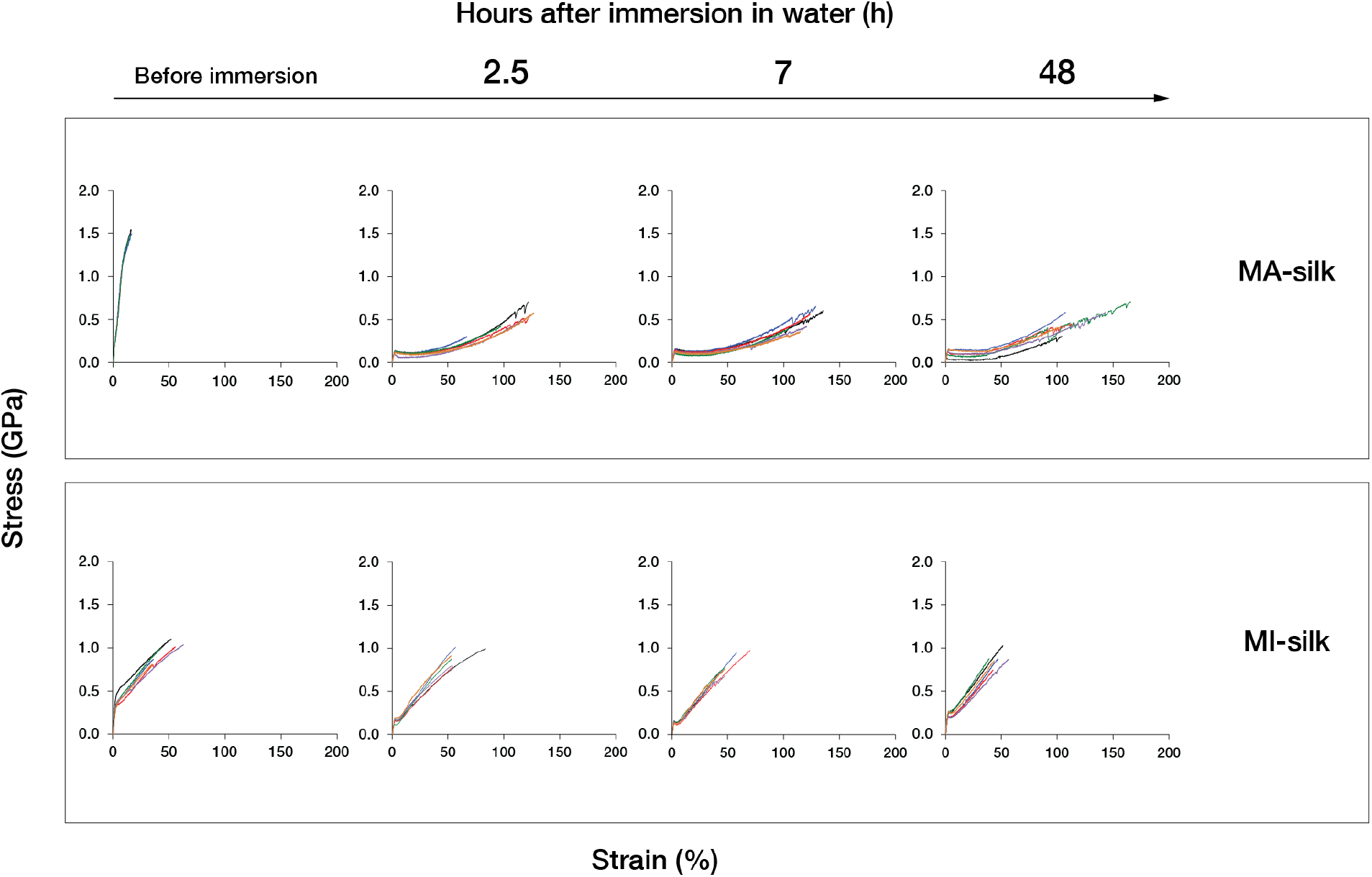
Stress‒strain curves of *A. ventricosus* MA-silk and MI-silk before and after immersion and subsequent drying

This difference in the stress‒strain curve before and after contraction has similar trends in the ratio of contraction summarized in Table 1. In both types of silks, there were clear yield point in stress-strain curve after water immersion which was not observed in the dried state. Diameter of the MA-silk clearly increased by supercontraction, but that of MI-silk remained unchanged. The mechanical properties tested are summarized in Table 2. Tensile strength and Young’s modulus decreased in MA-silk just after water immersion, and again, changes were minimal in MI-silk. Similarly, the birefringence of MA-silk was reduced by approximately 63% after wetting, agreeing with Work’s report of *A. diadematus* MA-silk,^27^ whereas that of MI-silk was reduced by only approximately 7%.

**Table 2.** Mechanical properties, diameter and birefringence of MA-silk and MI-silk before immersion in water and subsequent drying.

|  | Hours after immersion in water (h) | Tensile strength (GPa) | Strain at break (%) | Young's modulus (GPa) | Toughness (MJ * m-3) | Diameter (μm) | Birefringence (*10-3) |
| --- | --- | --- | --- | --- | --- | --- | --- |
| MA-silk | Before immersion | 1.406 ± 0.198 | 14.1 ± 3.9 | 13.62 ± 0.66 | 0.122 ± 0.052 | 3.40 ± 0.17 | 43.43 ± 2.92 |
|  | 2.5 | 0.509 ± 0.137 | 108.6 ± 22.8 | 5.27 ± 1.08 | 0.239 ± 0.082 | 4.83 ± 0.30 | - |
|  | 7 | 0.500 ± 0.124 | 121.1 ± 11.0 | 4.80 ± 1.38 | 0.277 ± 0.084 | 4.92 ± 0.23 | - |
|  | 48 | 0.510 ± 0.145 | 122.9 ± 25.8 | 5.67 ± 1.03 | 0.294 ± 0.140 | 5.02 ± 0.26 | 16.11 ± 0.98 |
| MI-silk | Before immersion | 0.970 ± 0.110 | 53.8 ± 10.6 | 13.58 ± 2.05 | 0.328 ± 0.101 | 2.11 ± 0.10 | 46.26 ± 1.13 |
|  | 2.5 | 0.898 ± 0.095 | 58.7 ± 12.1 | 8.86 ± 0.89 | 0.308 ± 0.098 | 2.45 ± 0.04 | - |
|  | 7 | 0.812 ± 0.121 | 51.91 ± 9.9 | 8.38 ± 0.58 | 0.234 ± 0.074 | 2.11 ± 0.13 | - |
|  | 48 | 0.879 ± 0.091 | 53.4 ± 9.3 | 11.44 ± 1.15 | 0.249 ± 0.057 | 2.14 ± 0.15 | 43.05 ± 1.50 |

### Proteome analysis of MA-silk and MI-silk from *A. ventricosus* and *T. clavata*

We analyzed the protein components of MA-silk and MI-silk reeled from *A. ventricosus* and *T. clavata* using nanoLC‒MS/MS and the iBAQ method. From *A. ventricosus* MA-silk, we detected major and minor ampullate spidroins and candidates of spider silk-constituting elements (SpiCEs) (Figure 3). The detected spidroins were MaSp1, MaSp2, MaSp3, MaSp4 and MiSp. Although the protein abundance varied among the samples, the sum of the fractions of MaSp1 and MaSp2 was consistently greater than those of MaSp3 and MaSp4. MiSp was the smallest amount of spidroin for all samples (≈(approximately 1%). Two genes of MaSp4 spidroin were detected from the proteome analysis, MaSp4A1.s1 (GBM28631.1) and MaSp4A1 (GBM28637.1) (Figure S3).

**Figure 3.**
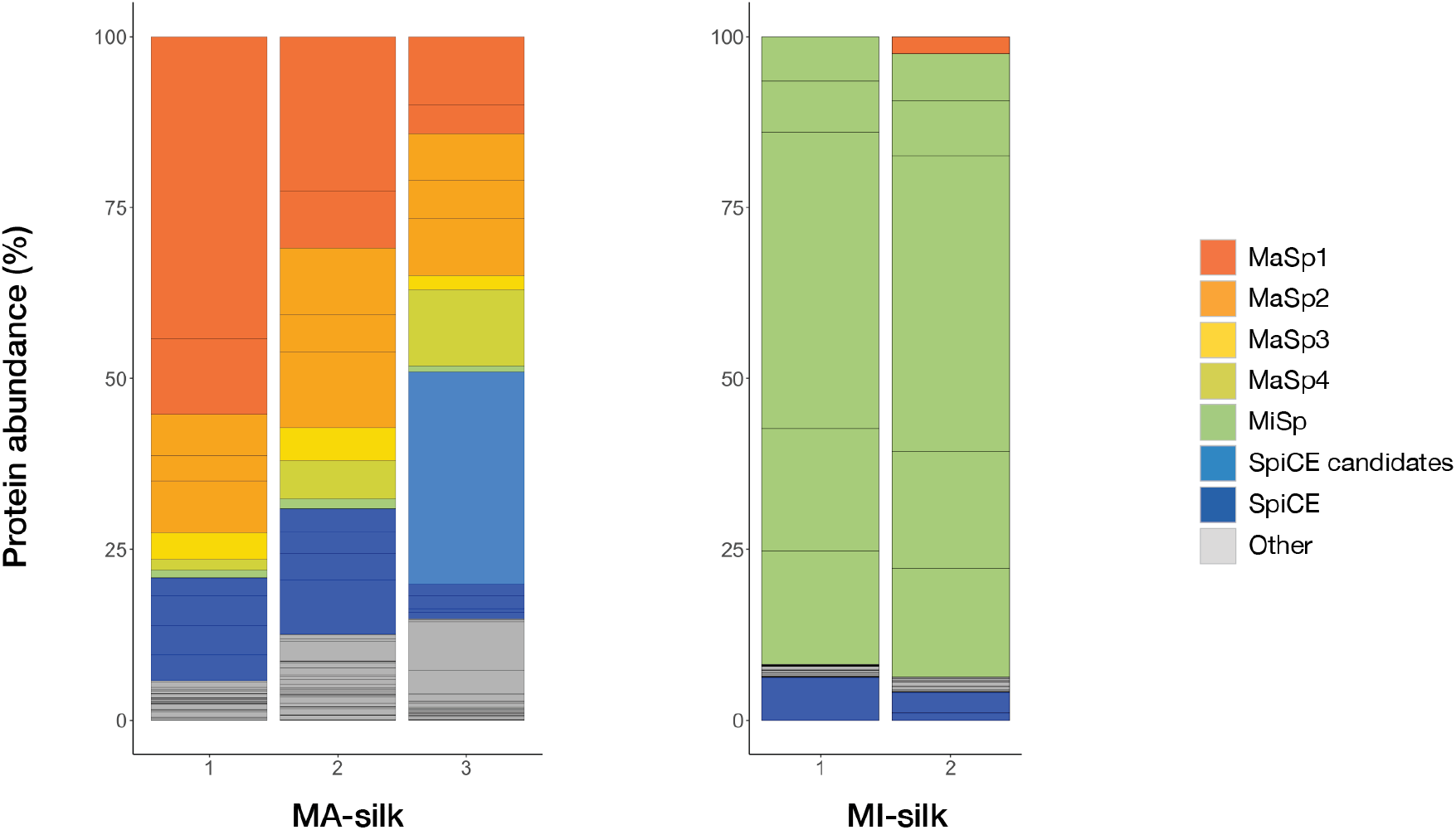
Protein abundance of *A. ventricosus* MA-silk and MI-silk

From *A. ventricosus* MI-silk, we detected major and minor ampullate spidroins and SpiCE candidates. The main component of MI-silk was 5 MiSps, and it also contained MaSp1. SpiCE candidate g6858.t1 was detected from all MA-silk and MI-silk samples (Figure S4), suggesting that the expression of SpiCE is common among the major and minor ampullate silks, contributing also to the mechanical properties of MI-silk, as suggested by Kono et al. for MA-silk.^16^ From *T. clavata* MA-silk, we detected MaSp1, MaSp2, MaSp3, SpiCEs and SpiCE candidates, and from MI-silk, MaSp1, MiSp and SpiCEs (Figure S5). Interestingly, the protein composition of MI-silk was strikingly different from that of *A. ventricosus*, where all samples contained a larger fraction of MaSp1 than MiSp. Similar to *A. ventricosus*, there were common SpiCEs contained both in MA-silk and MI-silk (Figure S6), of which SpiCE-NMa1 was previously shown to enhance the mechanical properties of MA-silk.^16^

### Fibers of recombinant spidroins

To compare mechanical properties and how each protein reacts against water immersion, we created recombinant mini-spidroins of MaSp1, MaSp2 and MiSp and created artificial spider silks. The constructs consisted of an N-terminal domain (NTD), a shortened repeat region and a C-terminal region, where the molecular weight was adjusted to approximately 50 kDa. For all constructs, lyophilized protein powder was dissolved in DMSO and 2 M LiCl at a concentration of 15% (w/w) and was coagulated with MeOH during subsequent drawing in a water bath. Spinning conditions were set according to a previous report.^16^ The maximum draw ratio was 6 x for MaSp1 and MiSp and 5.5 x for MaSp2. First, we compared the mechanical properties of artificial spider silks for each type of spidroin by tensile testing. Figure 4 shows stress-strain curves of 5 x drawing fiber for each recombinant mini-spidroin. Mechanical properties are shown in Table 3. MaSp1 had the highest tensile strength and the lowest strain at break. On the other hand, MaSp2 had the lowest tensile strength and the highest strain at break. Tensile strength and strain of MiSp was in between MaSp1 and MaSp2. Then, we tested the contraction of artificial spider silks by measuring the length before and after immersion in water by the same procedure with natural MA-silks and MI-silks. For 5 x drawing fiber (Figure S7), the ratio of contraction was 34.67 ± 1.15% for MaSp1, 41.33 ± 1.15% for MaSp2 and 29.33 ± 1.15% for MiSp after drying (Table 4). MaSp2 showed the largest contraction ratio, and MiSp showed the smallest contraction ratio. Although natural MI-silk showed approximately 7% contraction, artificial MiSp fiber showed an approximately 4 times larger contraction ratio than that. Additionally, while natural MA-silk showed approximately 50% contraction, the contraction ratios of artificial MaSp2 fiber and artificial MaSp1 fiber were lower than that. These differences between natural and artificial fibers were potentially due to differences in microstructural organization between natural and artificial silks, as contraction has been observed for regenerated silkworm silk fiber.^46^

**Table 3.**
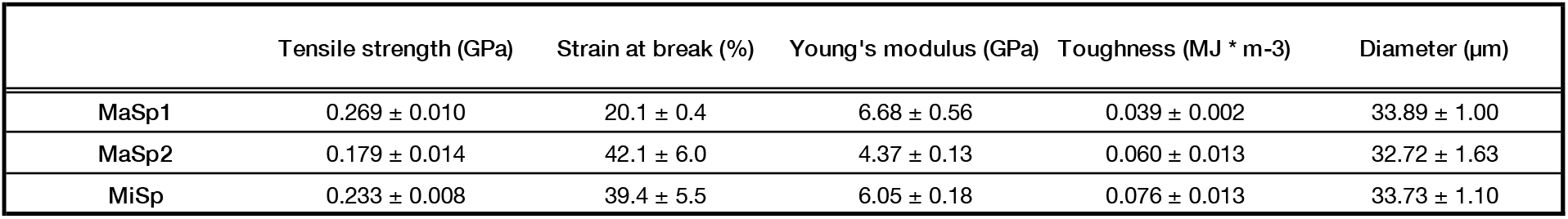
Mechanical properties and diameter of recombinant mini-spidroin fibers.

**Table 4.** Percentage of contraction for each recombinant mini-spidroins fibers.

|  | MaSp1 | MaSp2 | MiSp |
| --- | --- | --- | --- |
| % contraction by immersion in water | 32.67 ± 3.06 | 39.33 ± 1.15 | 26.67 ± 1.15 |
| % total contraction after drying | 34.67 ± 1.15 | 41.33 ± 1.15 | 29.33 ± 1.15 |

**Figure 4.**
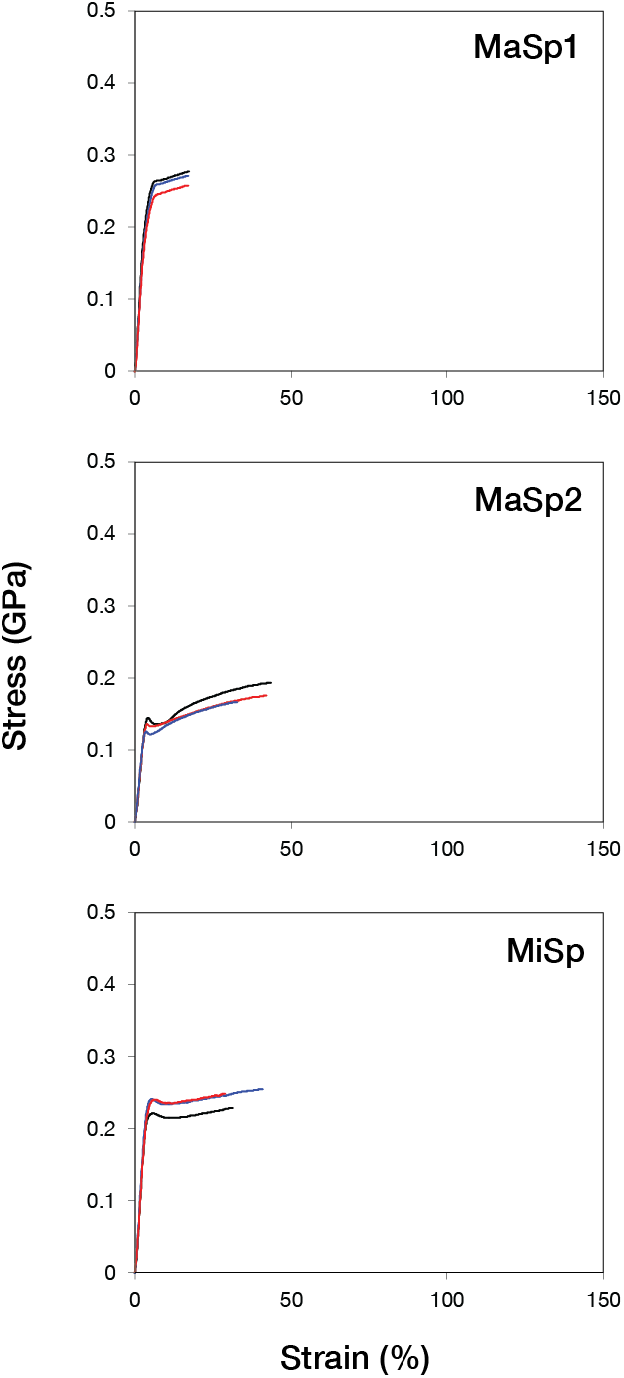
Stress‒strain curve of artificial fibers of recombinant MaSp1, MaSp2 and MiSp before and after water immersion

Tensile testing was also performed for the artificial spider silk samples after water immersion (Figure S8). After water immersion and subsequent drying, the strain at break of fibers of all mini-spidroins was increased. MaSp2 had the highest strain at break, and those of MaSp1 and MiSp were similar and approximately half of MaSp2. For tensile strength, MaSp2 was lower than MiSp and MaSp1. After water immersion, the stress‒strain curves of MaSp1 and MiSp showed similar trends.

## Discussion

Spider dragline silks are often composed of MA-silk and MI-silk, but the simultaneous analysis of detailed molecular differences between the two remains relatively less explored. Here, we studied the mechanical properties (Figure 1), microstructural organization (Figure S1 and Figure S2), effects of water immersion and subsequent drying (Figure 2 and Table 2) and protein composition of MA-silk and MI-silk (Figure 3 and Figure S5) that were separately silked. The mechanical properties of separately silked MA-silk and MI-silk from *A. ventricosus* were in line with previous reports,^2,28^ confirming the successful separate silking of two types of silks.

We then performed structural analysis by WAXS to observe nanostructural differences between MA-silk and MI-silk (Figure S1). The profile showed that both silks have semicrystallite structures, and the size of crystallites was different in silk types. With WAXS performed while drawing the fibers, we observed that MI-silk crystallites are more prone to deformation than MA-silk crystallites (Figure S2), supporting the aforementioned observation of irreversible changes in birefringence in MI-silk. The crystallites of MA-silk were larger than those of MI-silk on the *a*-axis and *b*-axis in *A. ventricosus*, which is different from the crystallite size comparison of *T. clavipes* MA-silk and MI-silk reported by Sampath et al., where MI-silk has been reported to have larger crystallites than MA-silk^31^ and suggests compositional differences between *Araneus* and *Trichonephila* MI-silks.

As previously reported, one of the most striking differences in the mechanical properties of these two fibers is their reaction to wetting.^27,28,31^ We obtained percentages of contraction of the MA-silk and MI-silk of *A. ventricosus* when immersed in water and with subsequent drying (Table 1). As expected, compared with the contraction ratio of *T. inaurata, A. trifasciata* and *A. diadematus* reported in previous studies, *A. ventricosus* MI-silk was more similar to those of *A. trifasciata* and *A. diadematus*. The difference in the contraction ratio between *Araneus* and *Trichonephila*, likewise the difference in the size of crystallites observed in structural analysis, suggest compositional difference in these genera.

While the stress-strain curve of MA-silk was significantly deformed by water immersion and resultedin increased strain at break and decreased tensile strength and Young’s modulus, MI-silk was less affected by water, as shown in Figure 2 and Table 2. On the other hand, an yield point in the stress‒strain curve became observable after wetting both in MA-silk and MI-silk. This indicates the loss of orientation in the micro-organizational structure in MI-silk upon wetting, as observed in the birefringence measurements before and after immersion in water. The decrease in birefringence could be an explanation for this change, mirroring the reports of a similar loss of orientation of the amorphous component in *Trichonephila clavipes* and *Argiope aurantia* MI-silk before and after immersion in water using WAXS by Sampath et al.^31^

Compositional analysis was thus conducted using proteome analysis of MA-silk and MI-silk of *A. ventricosus* and *T. clavata* (Figure 3 and Figure S5). MA-silk and MI-silk of both species were comprised of spidroins and minor components SpiCE, as previously reported,^16,17^ and interestingly, both species showed the inclusion of common SpiCEs both in MA-silk and MI-silk, suggesting that the use and enhancement of mechanical properties by SpiCE is a conserved feature among the major and minor ampullate silks. In particular, SpiCE-NMa1, a SpiCE previously shown to double the tensile strength of artificial spider silk just by 1% (w/w) addition, was included both in MA-silk and MI-silk of *T. clavata*. The sequence similarity of common SpiCEs of *A. ventricosus* and *T. clavata* was low (16% identity), confirming the clade-specific adaptation expected in our previous report.^17^ Notably, MI-silk was composed of MiSp and MaSp1 and not MaSp2. Moreover, the compositional ratio between MaSp1 and MiSp was strikingly different between *A. ventricosus* and *T. clavata*, where it was predominantly comprised of MiSp with little MaSp1 in *A. ventricosus* and a mix of these two with the excess of MaSp1 over MiSp in *T. clavata*. This reciprocal composition of MiSp and MaSp1 in the two species is in accordance with the reversed crystallite size among the two genera, and with the nanostructural organization in MA-silk, the larger crystallite size can therefore be attributed to MaSp1 being the major constituent. Unlike many other Orbiculariae, *Trichonephila* is known for leaving auxiliary spirals on its orb web. Hasselberg and Vollrath studied the mechanical properties of auxiliary spirals in the orb web of *Trichonephila edulis* and concluded that it was highly unlikely that all scaffolding *T. edulis* silk was MI-silk^23^. We argue that the MA-silk-like composition of *Trichonephila* MI-silk, with the major component being MaSp1 instead of MiSp, could explain this paradoxical use and mechanical property of MI-silk. The use of only MaSp1 among the MaSp paralogs in MI-silk both in *Araneus* and *Trichonephila* suggests that MaSp1 is the basic component for ampullate silks rather than MaSp2, MaSp3, and MaSp4, which is in good agreement with the suggestion that MaSp2 is relatively newly acquired at the base of Orbiculariae ^47,48^.

The proteomic compositional analysis suggests that the supercontraction and irreversible change in mechanical properties by water immersion of MA-silk can be attributed to the inclusion of MaSp2. The -GPG-motif shared in MaSp2, MaSp3 and MaSp4 is suggested to be the cause of supercontraction,^34,35^ whereas the mirroring -GXG-motif of MaSp1 was linked to Tg-related contraction, as proposed by Guan et al.^35^ To better characterize this contribution of different spidroins, we produced artificial spider silk made individually from recombinant mini-spidroins of MiSp, MaSp1 and MaSp2. As expected, artificial MaSp2 fiber was the most affected by water immersion and the percentage of contraction was the highest in these three spidroins, again supporting the fact that MI-silk is predominantly composed of MiSp and MaSp1 and has fewer effects against water immersion (Table 4).

## Conclusions

The difference in reaction to water immersion between MA-silk and MI-silk is an interesting model of silk component-structure‒function relationships. Here, we analyzed the difference in supercontraction behavior in two types of ampullate silk prepared by separate forcible silking. As previously observed, we confirmed that MI-silk has a lower ratio of contraction and a lower level of mechanical property deformation than MA-silk. We then performed proteome analysis of MA-silk and MI-silk enabled by the silking method. The MI-silk constituents were revealed by MaSp1, MiSp and SpiCE, and no MaSp2 was observed in their composition. Comparison of protein composition between *A. ventricosus* and *T. clavata* suggests that the difference in composition could explain the different ecological usage of MI-silk in the two species.

Furthermore, we created artificial fibers from recombinant MaSp1, MaSp2 and MiSp mini-spidroins and tested supercontraction. The fact that MI-silk does not contain MaSp2 and recombinant MaSp2 has a higher contraction ratio than MaSp1 and MiSp suggests that the reason for the difference in supercontraction is the existence of MaSp2.

## AUTHOR INFORMATION

### Author Contributions

N.K. and K.A. conceived and designed the research. H.N. performed silk sample preparation, tensile testing, birefringence measurement, and artificial silk spinning of recombinant mini-spidroins. K. N. and H.M. performed WAXS measurements and characterized the data. M.M. performed proteome analysis. N.K. prepared reference databases for proteome analysis. H.N. and K.A. wrote the manuscript. All authors reviewed the manuscript.

### Funding Sources

This work was supported by a grant from the ImPACT Program of the Council for Science, Technology and Innovation (Cabinet Office, Government of Japan), MEXT Program: Data Creation and Utilization-Type Material Research and Development Project Grant Number JPMXP1122714694, and by research funds from the Yamagata Prefectural Government and Tsuruoka City, Japan.

Notes

H.N. is an employee of Spiber Inc. The authors declare that they have no other competing interests.

## ACKNOWLEDGMENTS

We acknowledge T. Osawa, C. Sato, H. Sugawara, Y. Ito, H. Chi and the fermentation & purification team of Spiber Inc. for technical support in the construction of expression strains, fermentation, purification and spinning of mini-spidroins; and Y. Takai for technical support in proteome analysis.

---

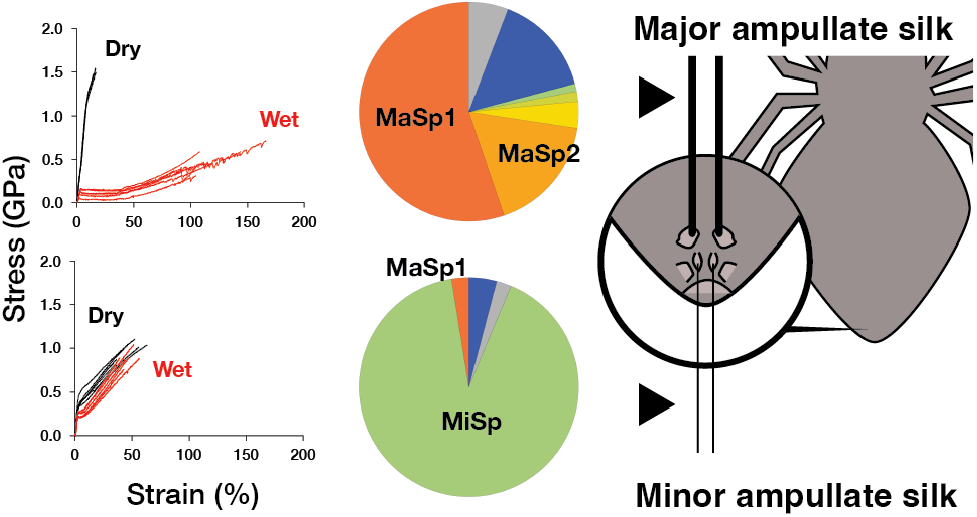

For Table of Contents Only

