## Supplementary material for "Composition of minor ampullate silk makes its properties different from those of major ampullate silk": Figure S

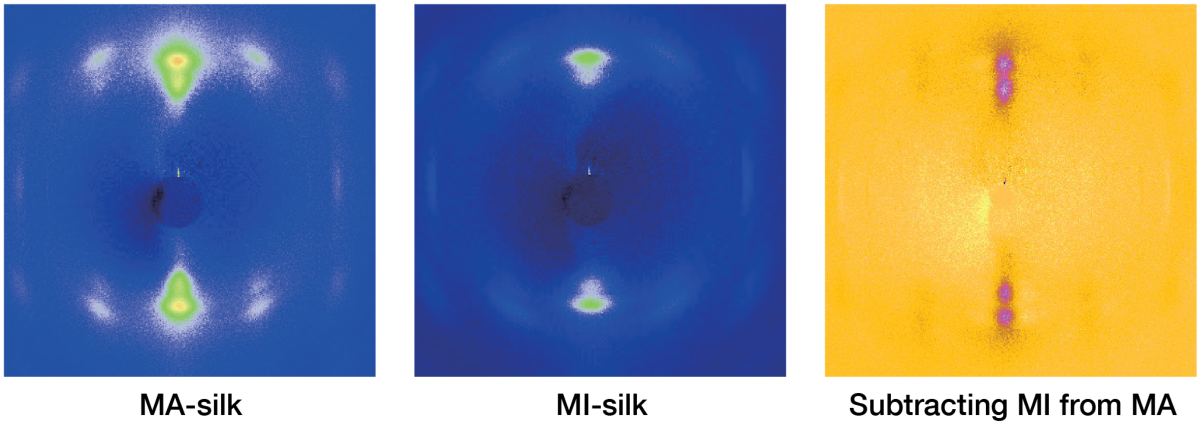


**Figure S1.** WAXS 2D profile of *A. ventricosus* MA-silk and MI-silk

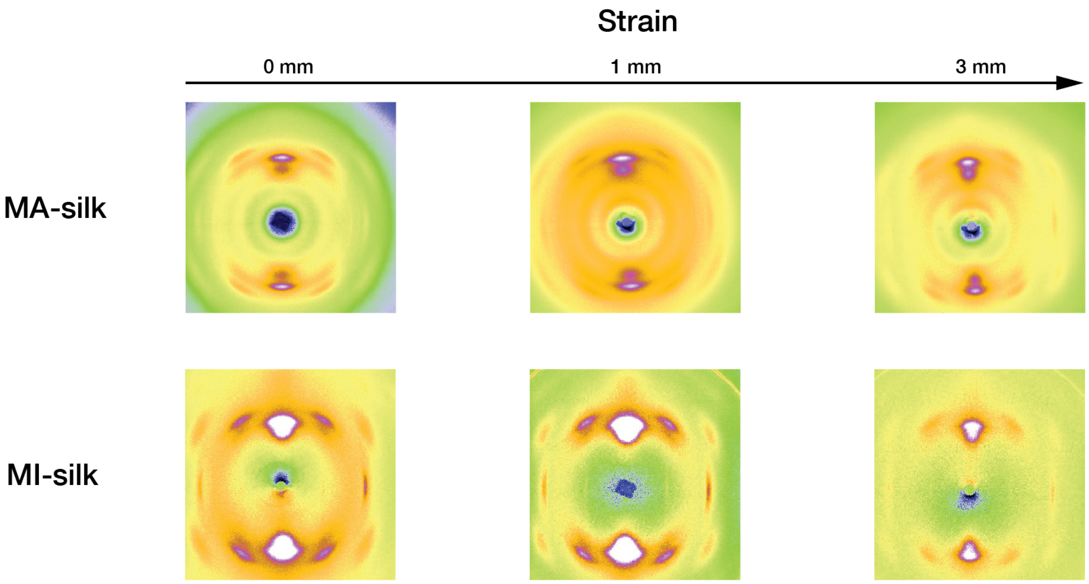


**Figure S2.** Structural analysis of *A. ventricosus* MA-silk and MI-silk upon extension

MaSp4A1.s1   GBM28631.1

```
MSWFKTFSLACLLVLCTQAVVVVEGARSPWESPQLAQSFINSFLRAISRSGAFSYSQLDD
MSTIGETLTIAMDKFTGSNKNIKSKLQALDMAFASSMAEIAVAEQGGLSISEKTNAIENA
LNAAFLETTGVIFTQFVSEIKSLIFLIAQASTNEISSMPTSGAGGYGQAASGPQGPGSSG
PSPQGPSGPTPQVPGPSSSVITSYGPGPQGPYGPGPQGPSPQGPTGPGPQGPGSSSSVIT
SYGPGPQGPYGPGPQGPSPQGPTGPGPQGPGSSSSVITSYGPGPQGPYGPGSQGPSPQGP
TGPGPQGPGSFSSVLSYGPGPQGPSGPGPQRPSPQGPTGPGPQGPGSSSSVISSGPGPQG
PSGPGPQRPSPQGPTGPGPQGPGSSSSVITSYGPGPQGPGSSSSVITSYGPAPQGPSGPG
PQGPSPQGPTGPGPQGPTLSSFTFSGPGPQGPSGPSPQRPSPQGPTGPGPQGPGSSSSVI
TSYGPGPQGPSGPGPQGPSPQGPTGPGPQGPGSSVSVLSYGPGPQGPSGPGPQGPSPQGP
SGPGPQGPGSSSSVITSYGPGPQGPYGPGPQGPSPQGPTGPGPQGPGS
```

MaSp4A1   GBM28637.1

```
MTLVLRSQAQWGLEVLGPGKQCTRNAKYRSPQGPTGPGPQGPTLSSFTFSGPGPQGPSGP
SPLRPSPQGPTGPGPQGPGSSSSVITSYGPGPQGPSGPGPQGPSPQGPTGPGPQGPGSSS
SVITSYGPGPQGPSGPGPQGPSPQGPTGPGPQGPSLSSFAFSGPGPQGPSGPSSQGPSSG
YGRGQQYGQSVVISSASSRLSSPSATSRISSAVSSLMTSGPRNPVGLSIALGNILSQIQS
SNPGLSGCESLVQALLEIASALIQILSVSSVGQVDFRAIGQSASIVGQALMQNLG
```

**Figure S3.** Peptide mapping of *A. ventricosus* MaSp4A1.s1 and MaSp4A1

g6858.t1    GBM55674.1

```
GTMQSVLLSVHIEIGGGPDTLVSQISADGKLSLKREDHYLSMDSLLLLTLTISLMFTGGQ
CYSESMSCSTVNGKTSCESQKSAGNVAGAASSARTGQGGSYQDASAMTGNTYTGQSASSR
TAMGNLPQGGFGPGPAPFAGGFAFPFGAPPTFAGGSGNPAGTSASDSANTGTGSTSFTGT
SGSDGTGAGGAFAGGFAGSGEFVGTAAGGFAGSGDSGVFVGGGFPLPFGVSAFGDGSSVP
SGTSASDSGNTGTGSTFVTSTSGSDGAGAGGFAGPSSGGFVGSAAGGFAGFGGSGVFVGG
GAIPGIPGFGTGGGVFVGNPAAMMGAGFPFNFVFGGGGFPFGRR
```

Peptides detected in MA-silk

```
GTMQSVLLSVHIEIGGGPDTLVSQISADGKLSLKREDHYLSMDSLLLLTLTISLMFTGGQ
CYSESMSCSTVNGKTSCESQKSAGNVAGAASSARTGQGGSYQDASAMTGNTYTGQSASSR
TAMGNLPQGGFGPGPAPFAGGFAFPFGAPPTFAGGSGNPAGTSASDSANTGTGSTSFTGT
SGSDGTGAGGAFAGGFAGSGEFVGTAAGGFAGSGDSGVFVGGGFPLPFGVSAFGDGSSVP
SGTSASDSGNTGTGSTFVTSTSGSDGAGAGGFAGPSSGGFVGSAAGGFAGFGGSGVFVGG
GAIPGIPGFGTGGGVFVGNPAAMMGAGFPFNFVFGGGGFPFGRR
```

Peptides detected in MI-silk

**Figure S4.** Peptide mapping of *A. ventricosus* g6858.t1 in MA-silk and MI-silk

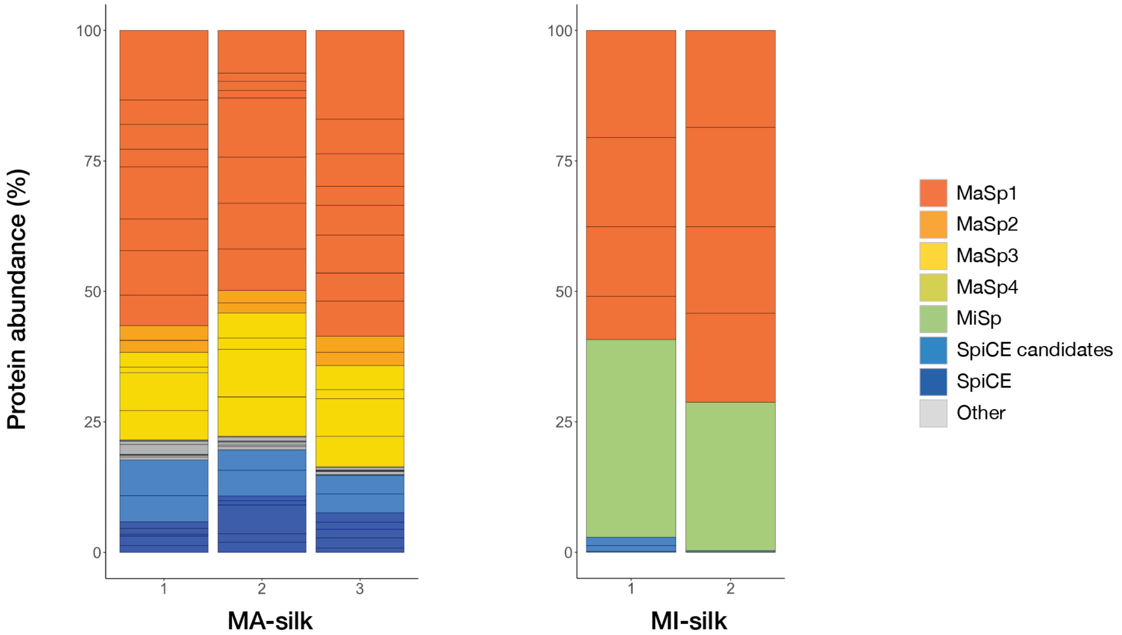


**Figure S5.** Protein abundance of *T. clavata* MA-silk and MI-silk

SpiCE-NMi1 GFQ84364.1

MNGGGLNGRTSGMNGGGLNGGGLNGGGLNGGGLNGGGFNGGGLNGGGFNGGNLKRGGFDQ
GGFGGLNGGGLNMGGFDDGGFNDGGLNMGGFDDGGFNDGGLNTGGLNSGGLRGGFKRGGL
NNGGLSGGGLSNGGFNGGNLDFGGFNDGGTNGGRLGSGNIGSGGNQGYDLGGWGPGGGFN
SGGNSLGGGMDISSGRGGGGGLNIPSRRGGGGGFNVPSGGGGLNIPSGGGGAGGGRGGGK
AVGVPSGGSGGSSGGWSFGNLFRGWNAGNAGGGNAIDLGSGGDSIGRGGRGGKKAAGGAA
GAASAASGAAATGNGGAGGAAGAASGAAAAGDGGSASAASGAAATGNGGAGGAAGAASGA
AAAGDGGSASAASGAAATENGGAGGEADAASGAASGAGANRSGGRARAAASAGASGNGGA
AGAGASSSAASGGGSRGGNSGGRCKAKGKDKQSDNGQDGDDDNGNMLDGLKGKLGNGIKG
KGQELRNKMDGDDLGRGMLGSLSGKRNSNSQSNNGNGNSAAASSWSYATENGDNSGDDDS
GNKGNGLFGMFWRRRNGGDQSGNGDGDDQSDNSGDDQPDNGDGDDQSGDDGSQDDSNTRG
NKLLGKLKSSNMMNGGGAPRNLDNKLLGKINRRRNGNNEGDSDSVNLDANQGGGGWGFGR
WFGQGNGGGIQGGRDISSSATSASSARAGDAGGIGQGRSFAVSESMANAGGRQGGSRAES
LSIANARGNGRRGSSAEATSMAQANGFSNSYDGSIGDQGNDNMMSAFLQFMKQKWIELNK
KGGNSVDIDSEWNKIVKGSGWLNGIGGSGNGLGMGSASGLGKGLGLGKGAGLGGASALGS
GSSNGIGLGGGQISDKGSKMKGFLKGLGGESDDDSGSNDDLGMDGDFGLNKVSRKKGLFG
GMGGGLGRGGGGGSGGGSGGGGGSGGGLGLGSGRGGGSGGGGLGSGGGGGSGRGGGSGGG
GGRGGGGGSGGGLGLGSGGGGGSGRGGGSGGGGGRGGGSGRGGGSGGGGGSGGGGGSGRG
GGSGLGGGSGLGGGSGLGGGSGGGGGSGGGGGLGGGGGLGSRSSNLGMDDGR

**Figure S6.** Peptide mapping of *T. clavata* SpiCE-NMi1

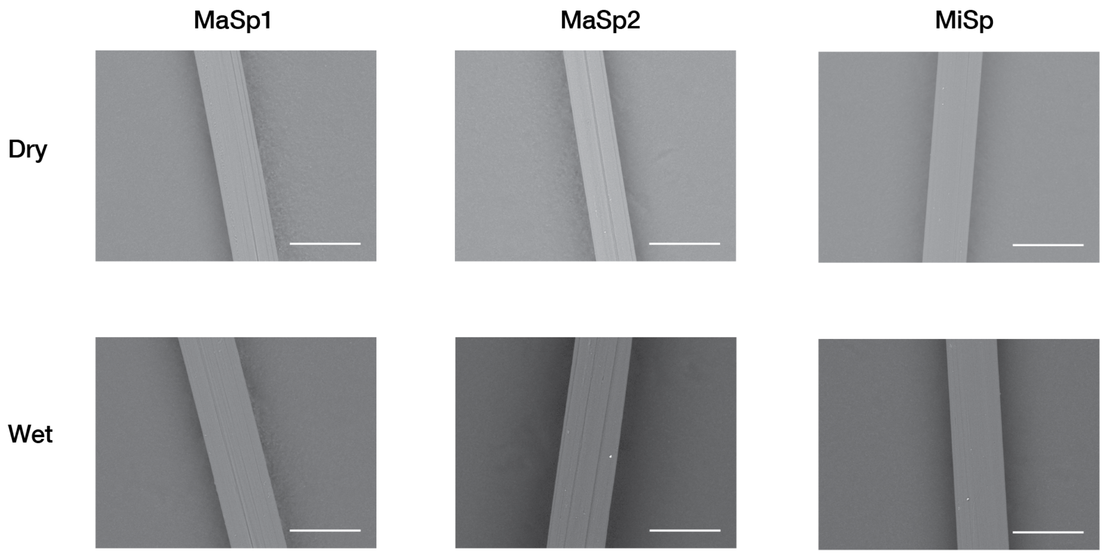


**Figure S7.** Morphology of artificial fiber of MaSp1, MaSp2 and MiSp, before and after water immersion

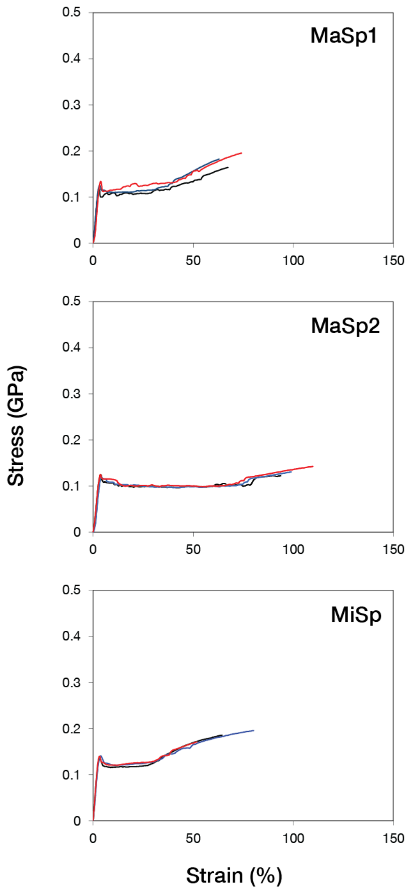


**Figure S8.** Stress–strain curve of contracted artificial fibers of recombinant MaSp1, MaSp2 and MiSp.

**Table S1.** Mechanical properties and diameter of recombinant mini-spidroin fibers after immersion in water and subsequent drying.

| | Tensile strength (GPa) | Strain at break (%) | Young's modulus (GPa) | Toughness (MJ * m-3) | Diameter (μm) |
|---|---|---|---|---|---|
| **MaSp1** | 0.180 ± 0.016 | 70.9 ± 4.5 | 4.54 ± 0.42 | 0.089 ± 0.013 | 42.09 ± 1.65 |
| **MaSp2** | 0.132 ± 0.009 | 102.8 ± 8.1 | 3.75 ± 0.57 | 0.107 ± 0.013 | 40.03 ± 2.04 |
| **MiSp** | 0.184 ± 0.012 | 68.1 ± 12.8 | 5.02 ± 0.19 | 0.093 ± 0.026 | 38.25 ± 0.58 |
